# Simple amplicon sequencing library preparation for plant root microbial community profiling

**DOI:** 10.1101/2021.04.14.439905

**Authors:** Kie Kumaishi, Erika Usui, Kenta Suzuki, Shungo Kobori, Takumi Sato, Yusuke Toda, Hideki Takanashi, Satoshi Shinozaki, Munehiro Noda, Akiko Takakura, Kayoko Matsumoto, Yuji Yamasaki, Hisashi Tsujimoto, Hiroyoshi Iwata, Yasunori Ichihashi

## Abstract

Microbiota are a major component of agroecosystems. Root microbiota, which inhabit the inside and surface of plant roots, play a significant role in plant growth and health. As next-generation sequencing technology allows the capture of microbial profiles without culturing the microbes, profiling of plant microbiota has become a staple tool in plant science and agriculture. Here, we have developed a novel high-throughput method based on a two-step PCR amplification protocol, involving DNA extraction using magnetic beads and PCR purification using exonuclease, for 16S rRNA gene amplicon sequencing of plant root microbiota. This method reduces sample handling and captures microbial diversity comparable to that obtained by the standard method. We found that using a buffer with magnetic beads enabled efficient extraction of microbial DNA directly from plant roots. In addition, we demonstrated that purification using exonuclease before the second PCR step enabled the capture of higher degrees of microbial diversity, thus allowing for the detection of minor bacteria compared with the purification using magnetic beads in this step. Our method offers a simple and high-throughput solution for maintaining the quality of plant root microbial community profiling.

## Background

The plant root is a key underground organ that interacts with soil, in which one of the richest microbial ecosystems on Earth exists [1]. Plant root microbiota, which inhabit the inside and surface of plant roots, improve plant growth by producing phytohormones, supplying nutrients, and protecting plants against pathogens and environmental perturbations, including drought and climate-dependent salinity changes [2–7].

Extensive efforts by the Earth Microbiome Project have led to the characterization of global taxonomic and functional microbial diversity, including the microbiome associated with plants [8]. In addition, many studies targeting plant microbiomes have been carried out, leading to the accumulation of large datasets [1,9–12], which will be utilized in agricultural applications through industry-academic collaborative projects [13]. Given that, along with the microbiome data, multi-omics analysis has been utilized in agricultural studies [14,15], high-throughput methods for the detection and analysis of plant microbiomes have become increasingly necessary.

Next-generation sequencing technology is continuously improving platforms to increase sequencing speed and quality. Even as sequencing capacity increases, sample library preparation is still laborious and time-consuming, which is a limiting factor for upgrading and expanding plant microbiome databases. Currently, many plant microbiome-based studies have used a kit-based or traditional DNA extraction method in a two-step PCR amplification protocol employed in microbiome amplicon sequencing (amplicon-seq) [12,16–19]. For example, the standard method (using isopropanol) requires 19 steps and takes approximately 2 h for DNA extraction, whereas 8 steps and approximately 30 min are required for PCR purification in library preparation (using AMPure XP beads) (**Fig. 1**). In this study, we present several improvements to the standard protocol of 16S rRNA gene amplicon-seq that have ultimately resulted in a method that requires 8 steps (30 min) for DNA extraction and 2 steps (10 min) for PCR purification in library preparation. Thus, we have developed a simple option that can be easily integrated into the automated process.

**Fig. 1.**
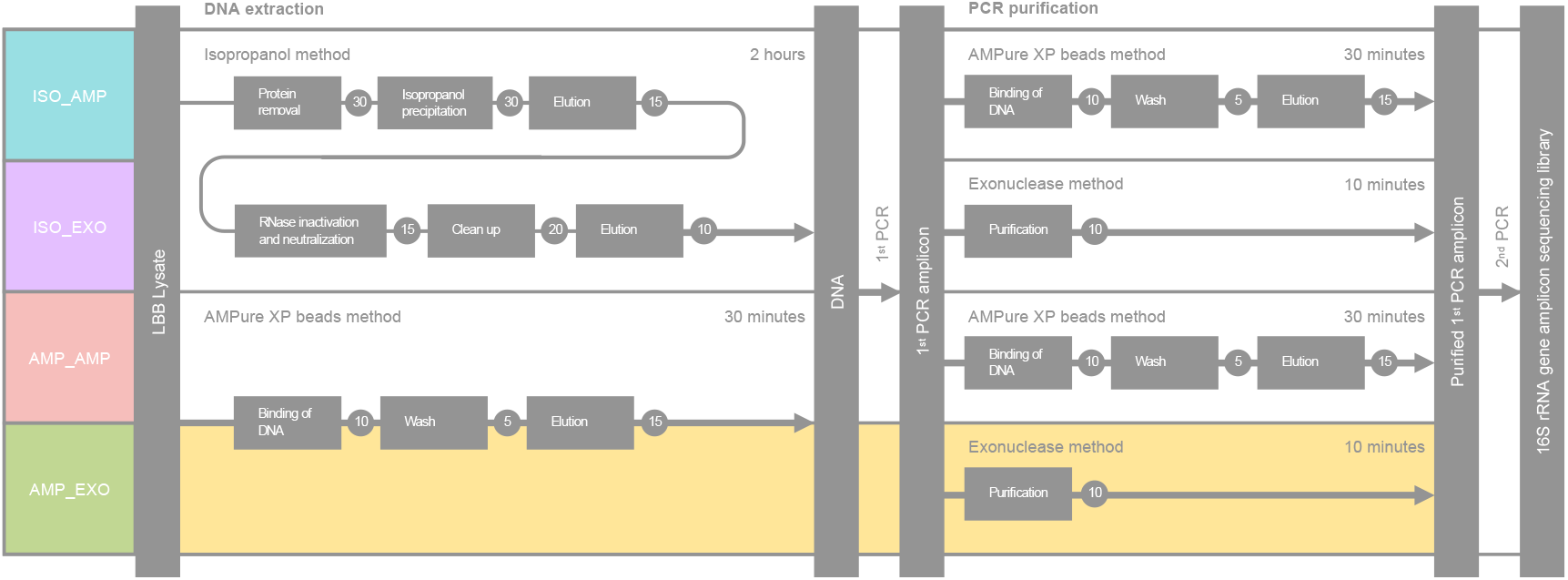
Schematic overview of the methods tested in this study. Experimental procedure for microbiome analysis. For DNA extraction, the AMPure XP bead method and isopropanol method as standard methods were tested. For the first PCR purification in the library preparation, the exonuclease and AMPure XP bead methods were tested.

## Results

### Yield and quality of DNA extraction among different methods

We initially searched for commercial DNA extraction methods for plant tissue and soil samples to develop a high-throughput experimental procedure; however, we ultimately found that utilizing AMPure XP beads combined with the proper buffer enabled the extraction of DNA from plant tissue and soil samples. When we compared the use of Tris-EDTA (TE) buffer and Lysate Binding Buffer (LBB) in the process of magnetic bead binding, DNA was successfully extracted from plant roots and soil samples when using LBB rather than TE (**Additional file 1: Figure S1**). In addition, the AMPure XP beads method using LBB showed relatively higher (though not statistically significant) yields of extracted nucleic acids (ng/100 mg powdered sample) than those obtained with the standard isopropanol method [20,21] (**Fig. 2a**). Notably, unlike the standard isopropanol method, the AMPure XP beads method does not include RNase treatment, which could be a reason for the resultant higher yield of extracted nucleic acids. On the other hand, both methods showed a 260/280 absorbance ratio of ∼1.8, suggesting that the extracted nucleic acids were relatively pure DNA from the plant tissue sample (**Fig. 2b**). Similar to the isopropanol method, the AMPure XP beads method showed high molecular weight DNA bands in agarose gel electrophoresis analysis (**Additional file 1: Figure S1**). Given that the AMPure XP beads method achieved a 75% reduction in the sample handling time compared to the isopropanol method (**Fig. 1**), the DNA extraction using AMPure XP beads could prove to be a high-throughput option that maintains a compatible yield and quality with that of the standard isopropanol method for plant roots and soil samples.

**Fig. 2.**
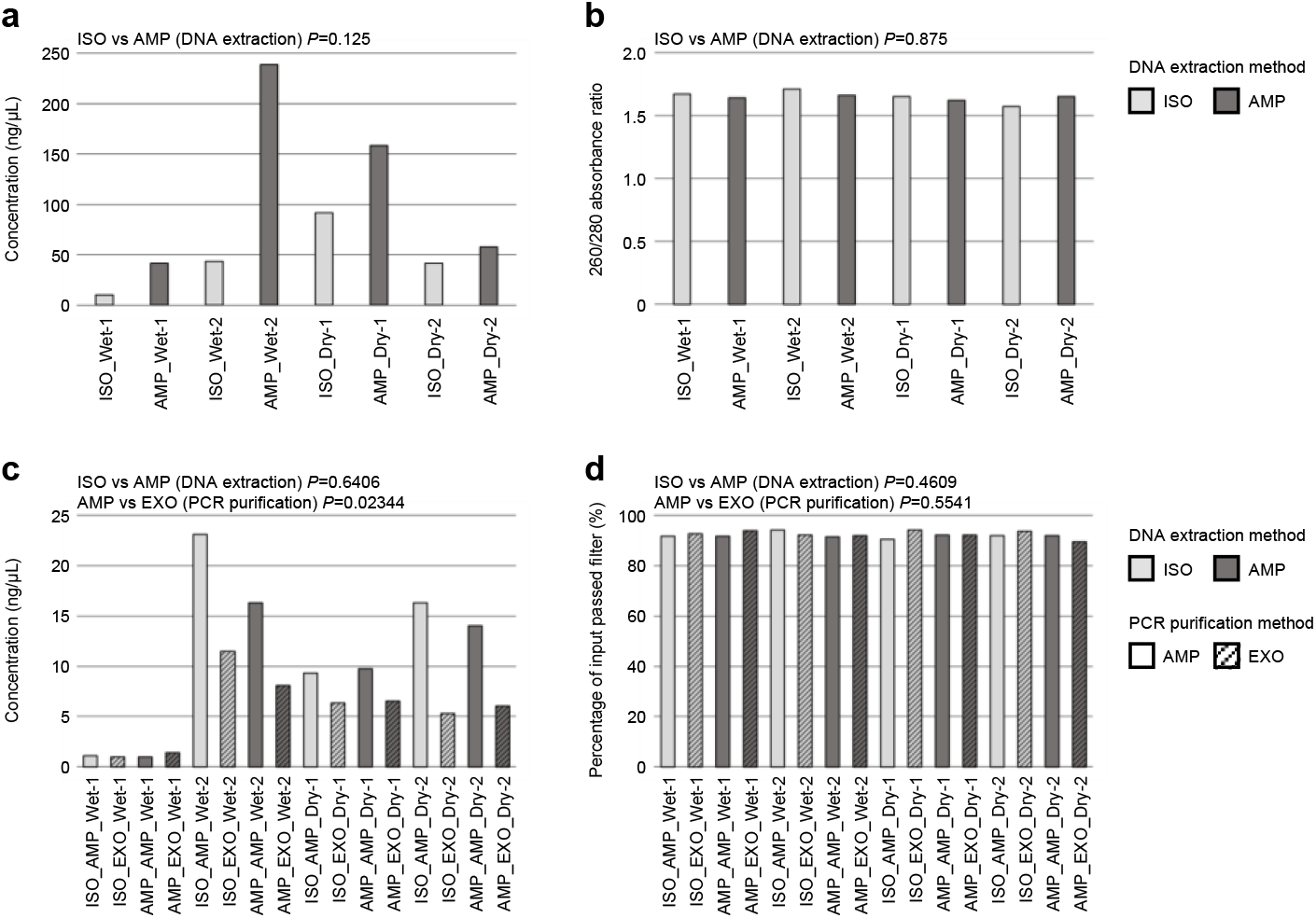
Yield and quality of DNA extraction and library preparation among different methods. The yield (ng/100 mg sample tissue) (**a**) and quality (260/280 absorbance ratio) of nucleic acid (**b**) are shown for the AMPure XP bead and isopropanol methods. Yield (ng/µL) (**c**) and reads passing set quality filter (%) (**d**) are shown for all methods described in Figure 1.

### Yield and quality of library preparation among different methods

The 2-step PCR amplification protocol is a common method used for microbiome amplicon-seq. During library preparation, exonuclease treatment is used for purification of the products obtained from the first PCR [14,22]. To assess the effect of the exonuclease treatment combined with our DNA extraction protocol, we compared the methods with all combinations of extraction and purification protocols, that is, the isopropanol method for DNA extraction and AMPure XP beads method for PCR purification (ISO_AMP), isopropanol method for DNA extraction and exonuclease method for PCR purification (ISO_EXO), AMPure XP beads method for DNA extraction and AMPure XP beads method for PCR purification (AMP_AMP), and AMPure XP beads method for DNA extraction and exonuclease method for PCR purification (AMP_EXO) (**Fig. 1**). Since we used DNA solutions with equal concentrations for 16S rRNA gene amplicon-seq library preparation, the concentration of the library was a result of the differences in the yield of nucleic acids among different DNA extraction methods (**Fig. 2c**). The exonuclease method for PCR purification in the library preparation showed a lower yield compared to the AMPure XP beads method for PCR purification (**Fig. 2c**). The AMPure XP beads method for PCR purification also performs size selection, such that the size selection before the second PCR (prior to final size selection) might enrich the first PCR product with the target size, leading to a high yield of library products. In contrast, size selection before the second PCR is associated with a risk of biased generation of specific amplicons; the exonuclease method could reduce this risk and rescue the minor PCR amplicons. Sequencing of the V4 region of bacterial 16S rRNA gene was carried out using 16 samples, which included all combinations of the methods that we tested in this study. A total of 120,849 reads with a mean read count of 7,553 reads per sample and a range of 1,510–24,782 reads were obtained. A total of 89.58-94.15% of reads passed the set quality filter (**Fig. 2d**). Notably, the exonuclease method, in addition to maintaining the sequencing quality, achieved greater than 60% reduction in the sample handling time compared to that required for the AMPure XP beads method (**Fig. 1**).

### Comparison of the diversity of plant microbial community observed among different methods

The alpha diversity based on the number of observed species, such as ASV, Shannon index, and Faith phylogenetic diversity, were compared among the methods. The exonuclease method detected ∼96% more ASVs (*P* < 0.05, **Fig. 3a**) and significantly increased the alpha diversity relative to the AMPure XP beads method for PCR purification (*P* < 0.05, **Fig. 3b and c**). These data support the idea that the exonuclease method could rescue the minor PCR amplicons, suggesting that this method can capture higher degrees of microbial diversity. To evaluate these methods based on the profiling efficiency of the microbial communities, principal coordinate analysis (PCoA) based on Bray-Curtis distances and weighted UniFrac distances [23] was performed. As a result, samples with two water conditions and two biological replicates were separated from each other, while samples processed using different methods were clustered together in the PCoA space considering both Bray-Curtis distances and weighted UniFrac distances (**Fig. 3d and e**). This indicates that the effect of differences among methods was much smaller for the microbial community profile than that of the biological sample differences, suggesting that our modified method (AMP_EXO) has a similar ability to capture the overall profile of the microbial community as that obtained using the standard method.

**Fig. 3.**
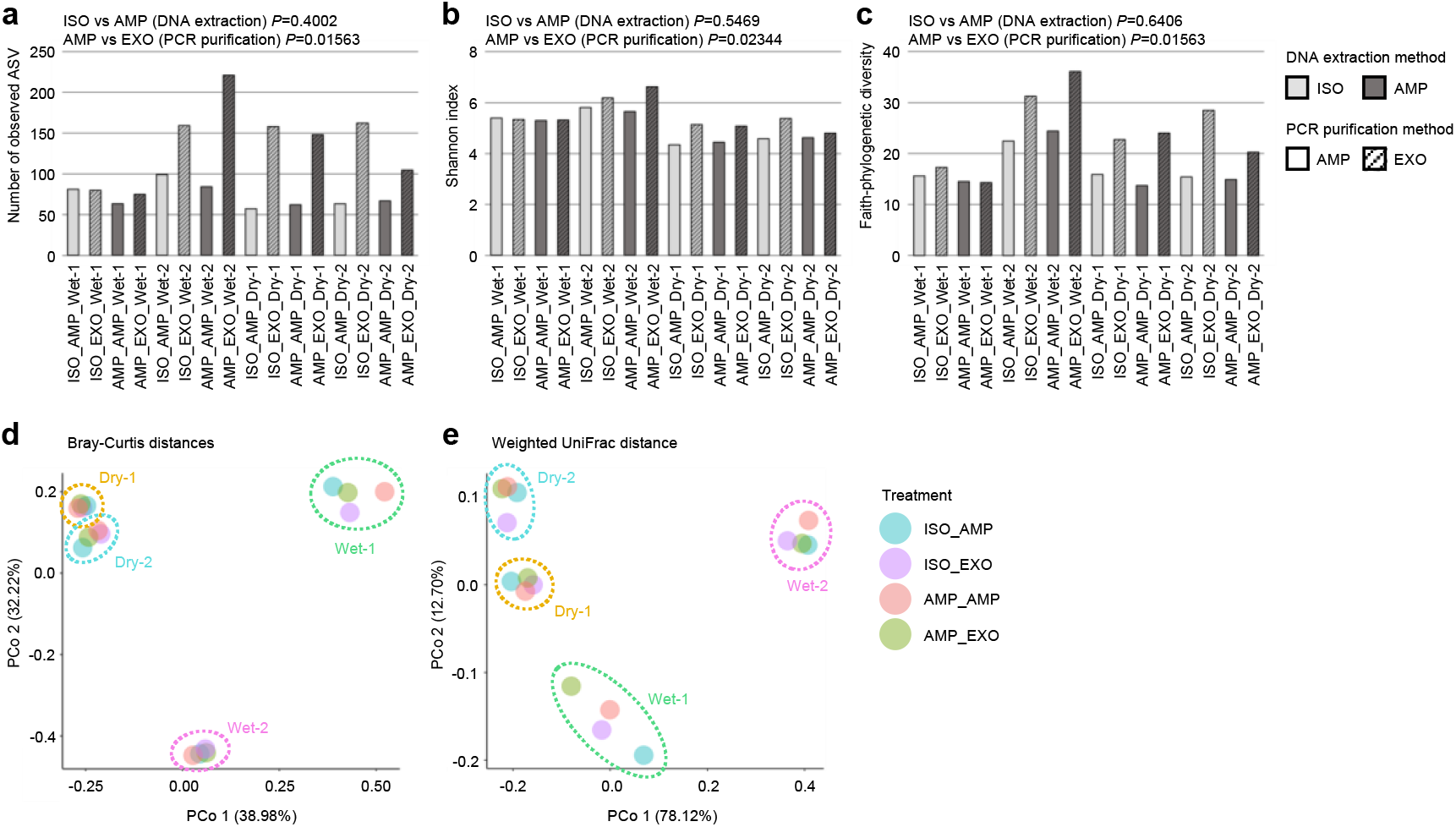
Plant microbial community diversity among different methods. Alpha diversity metrics of the number of observed ASVs (**a**), Shannon diversity (**b**), and Faith phylogenetic diversity (**c**). Principal coordinate analysis based on Bray-Curtis distances (**d**) and weighted UniFrac distances (**e**). The circles with different colors indicate different biological samples.

### Comparison of the taxonomic profile of plant root microbiome obtained using different methods

Our data showed that *Actinobacteria, Proteobacteria, Firmicutes, and Bacteroidetes* were the most abundant bacterial phyla, while *Streptomycetaceae, Oxalobacteraceae, Burkholderiaceae, Bacillaceae, Chitinophagaceae, Paenibacillaceae*, and *Xanthobacteraceae* were the most abundant bacterial families in our samples (**Fig. 4a and b**). The taxonomic profile detected order *Rhizobiales*, which includes *rhizobia* [24] and is similar to that obtained from soybean rhizosphere soil samples reported in previous studies [25,26]. In addition, our data showed that the phyla *Actinobacteria* and *Firmicutes* were enriched and *Proteobacteria* and *Bacteroidetes* were depleted in the root microbiota of soybeans cultivated under water-limited drought conditions (**Fig. 4a and b**); this is consistent with the results of a previous study that investigated the rhizosphere communities of several plant species [27–29], confirming that our method can generate comparable microbiome profile data. Furthermore, we detected no significant differences in the abundance of the gram-positive *Bacillus* sp. among the different methods (*P* > 0.1, **Fig. 4c**), showing that the DNA extraction method using AMPure XP beads has the ability to extract DNA from gram-positive bacteria with a thick peptidoglycan layer of the cell wall, which is comparable to that of the standard methods, such as the isopropanol method. Hierarchical clustering heat map based on the 20 most abundant ASVs showed that samples clustered into four groups according to the different biological samples and not according to the different methods (**Fig. 4d**). Furthermore, our LEfSe analysis discriminating four different methods showed no significant differences (*P* > 0.05); nonetheless, the following varied minor bacteria were identified based on different DNA extraction and PCR purification methods (*P* < 0.05): *Enterobacteriaceae* (average 2.46% relative abundance, enriched in ISO) and *Planctomycetales* (0.07%, enriched in AMP) for the tested DNA extraction method; *Xanthobacteraceae* (0.25%, enriched in EXO), *Sphingobacteriaceae* (0.21%, enriched in EXO), *KD4_96* (0.08%, enriched in EXO), *Noviherbaspirillum* (0.07%, enriched in EXO), *Acetobacterales* (0.06%, enriched in EXO), *Nitrospirota* (0.05%, enriched in EXO), and *Bacteriovoracales* (0.03%, enriched in EXO) for the tested PCR purification method. Thus, our modified method (AMP_EXO) can capture similarly abundant taxonomic profiles of plant root microbiome when compared to those obtained using standard methods, as well as show the potential to detect minor bacteria (less than 0.3% relative abundance, **Additional file 1: Figure S2**).

**Fig. 4.**
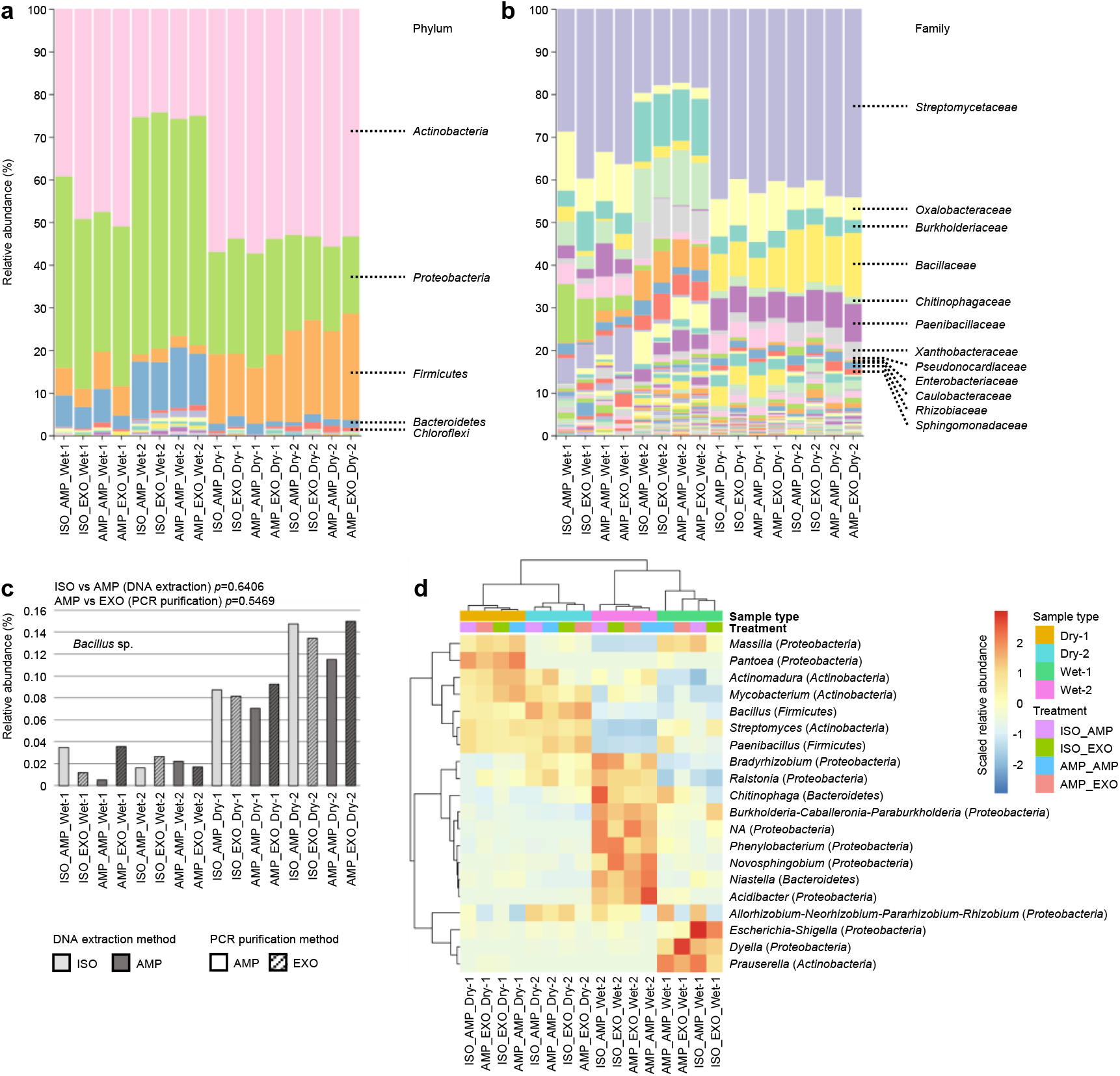
Taxonomic profile of plant root microbiome among different methods. The relative abundances of major phyla (**a**), family (**b**), and gram-positive *Bacillus* sp. (**c**) are shown. (**d**) Hierarchical clustering heat map based on the relative abundance of the 20 most abundant ASVs.

## Discussion

We have developed a new, simple, and high-throughput amplicon-seq library preparation method for plant root microbial community profiling, providing benchmarking data comparing between the newly developed method and other standard methods (**Figs. 1-3**). In contrast to the standard protocols, our method generated high-quality plant root microbiome data with a marked improvement in the ability to detect minor bacteria (**Fig. 3**). The data showed good agreement with that showing the taxonomic profile of the soybean rhizosphere microbiome, as well as successfully detected the changes in the taxonomic profile of the rhizosphere in response to drought treatment, a phenomenon also reported in previous studies [27–29] (**Fig. 4**). Our method uses magnetic beads for DNA extraction and exonuclease treatment for PCR purification, both of which are compatible with an automated process; this is expected to reduce sample handling and human errors compared to other protocols, thus enabling the simultaneous sequencing of thousands of samples.

Previous studies regarding plant microbiomes have commonly used column-based methods for DNA extraction, including bead-beating to lyse bacterial cells in plant tissues [1,10,12,16–18,30–37] and soils [38–43]. The column-based method requires laborious and time-consuming procedures, as well as a large amount of sample (> 100 mg). Our method using magnetic beads has successfully minimized the number of procedures to enable the extraction of DNA from low amounts of sample (∼20 mg), which is reflected in the fact that the reaction is completed in a single tube (**Fig. 1**). Although single-step PCR for library preparation has been previously used for plant microbiome studies [1,10,30–35,37,38,40,43], two-step amplification avoids potential bias due to the use of different indices in each primary amplification [39]. The standard method, including the method provided by Illumina, used AMPure XP beads for PCR purification before the second PCR, and a custom protocol used for phyllosphere microbiome study used exonuclease treatment [19]; however, no study compared these different methods employed for PCR purification before the second PCR.

One of the achievements of this study is finding a buffer that can be utilized for DNA extraction using magnetic beads. The buffer LBB was originally developed for RNA-seq library preparation in our previous study and can be used for DNA extraction with the isopropanol method [20,21]. During the search for a commercial DNA extraction method to develop high-throughput experimental procedures, we found that AMPure XP beads combined with LBB buffer showed a good yield of extracted DNA from plant tissues (**Fig. 2**). Since we have applied our DNA extraction method not only for plant root samples, but also for soil samples (**Additional file 1: Figure S1**), this method can be further applied for microbiome analysis of environmental samples; nonetheless, an assessment and fine-tuning of the method for various types of soils is necessary. In addition, recent studies pertaining to plant microbiomes have been focused on the functional aspects of microbiota at the gene level using metagenome sequencing in addition to taxonomic community profiling [11]; our method of DNA extraction using magnetic beads can be applied to fulfill the demand of high-throughput options for metagenome sequencing.

Another significant finding of this study is that the exonuclease treatment for PCR purification showed a high ability to capture higher degrees of microbial diversity, especially minor bacteria (**Figs. 3 and Additional file 1: Figure S2**). Rare bacterial species are increasingly recognized as crucial components of Earth’s ecosystems [44]. Several studies have shown that low-abundance plant-associated microbes enhance crop productivity and defense [45–47]. Given that our method can detect minor bacteria and capture the abundant taxonomic profile (**Figs. 3 and 4, Additional file 1: Figure S2**), this methodology would certainly contribute to the systematic accumulation of high-quality microbiome data.

## Conclusions

We have successfully developed a simple and high-throughput amplicon-seq library preparation method for plant root microbial community profiling. Using this method, we have produced libraries not only from soybean, but also from *Oryza* sp., *Brachypodium* sp., and *Brassica* sp., in addition to producing more than 1,000 libraries from plant roots cultivated in different agricultural fields from gray lowland soil to andosol (Ichihashi lab, unpublished results). Our method with reduced sample handling and compatibility with automated processes will be instrumental in future microbiome research with large-scale data. Although we have worked with limited samples, the method could be easily modified to target a broad range of environmental samples, such as those from various soils.

## Methods

### 1. Sample collection

Soybean (*Glycine max* (L.) Merr., Peking) was sown in a sandy field at the Arid Land Research Center, Tottori University, Japan, in July 2018. Plants were cultivated under two water conditions (well-watered and water-limited drought block) in two replicates. White mulching sheets (Tyvek, Dupont, US) and watering tubes were installed to control the soil conditions. Artificial irrigation from watering tubes was applied for 5 h daily in the well-watered blocks, while no artificial irrigation was used in the water-limited blocks from 14 days after sowing. Sixty-two days after sowing, the plant roots were harvested and washed with tap water. The tips (∼2 cm in length) of lateral roots developed from the main roots at 0-10 cm from the shoot/root junction were collected and kept at -20°C until sample preparation. The sampled root tissues were thought to contain endophytes and may have also contained the bacteria remaining from the rhizoplane.

### 2. DNA extraction

The collected root tissues were ground to a fine powder using a Multi-Beads Shocker (MB2200(S), Yasui Kikai Co. Osaka, Japan). For each of the collected tissue samples, 500 mg of the powdered sample was transferred into a 1.5 mL tube cooled by liquid nitrogen. One mL of lysate binding buffer (1 M LiCl Sigma-Aldrich, Cat. #L7026-500ML; 100 mM Tris-HCl, Wako, Cat. #318-90225; 1% SDS, Wako, Cat. #313-90275; 10 mM EDTA pH 8.0, Wako, Cat. #311-90075; Antifoam A, Sigma-Aldrich, Cat. #A5633-25G; 5 mM DTT, Wako, Cat. #048-29224; 11.2 M 3-Mercapto-1,2-propanediol, Wako, Cat. #139-16452; DNase/RNase-free H_2_O, Thermo Fisher Scientific, Cat. #10977015) [20] was added to the sample, which was then homogenized by vortexing, followed by incubation at room temperature (∼22°C) for 5 min. The tube was centrifuged at 15,000 rpm for 10 min at room temperature, and the supernatant (LBB lysate) was transferred to a new 1.5 mL tube. DNA extraction was performed using the following two methods: our custom protocol involving isopropanol extraction method [20] as the standard method, and extraction using AMPure XP beads (Beckman Coulter, Cat. #A63881). DNA concentration and absorbance were measured with a spectrophotometer (NanoDrop One^C^ Microvolume UV-Vis Spectrophotometer with WiFi, Thermo Fisher Scientific, Cat. #ND-ONEC-W).

#### 2-1. Isopropanol method for DNA extraction

The detailed method has been described in our previous publications [20,21] and was also used for soil and plant root microbial community profiling [14]. Briefly, LBB lysate (200 μL) was added to a 1.5 mL tube, and 5 μL of 10 mg/mL proteinase K was added to it; the mixture was then incubated at 37°C for 30 min. Next, 200 μL of 100% isopropanol was added to this sample, and this mixture was mixed gently, incubated at room temperature for 5 min, and centrifuged at 15,000 rpm for 5 min. The supernatant was discarded to avoid pellet loss, and 400 μL of 100% acetone was added to the tube. This was mixed gently, incubated at room temperature for 5 min, and centrifuged at 15,000 rpm for 5 min. The supernatant was carefully discarded to avoid pellet loss, and the pellet was dried. After repeated decolorization using acetone, we added to the tube 100 μL of 10 mM Tris-HCl (pH 7.5), incubated at 65°C for 10 min, and subsequently centrifuged at 15,000 rpm for 1 min. The supernatant was transferred to a new 1.5 mL tube, and 1 μL of 1 μg/μL RNase A was added and incubated at 37°C for 15 min. The supernatant was added to 10 μL of 3 M ammonium acetate and 250 μL of 100% ethanol, mixed, incubated at room temperature for 5 min, and centrifuged. The supernatant was discarded, and 400 μL of 80% ethanol was added, mixed, incubated at room temperature for 2 min, and centrifuged at 15,000 rpm for 1 min. The supernatant was carefully discarded to avoid pellet loss, and the pellet was dried. DNA was eluted in 50 μL of 10 mM Tris-HCl (pH 7.5).

#### 2-2. AMPure XP beads method for DNA extraction

Fifty μL of LBB lysate was put into 1.5 mL tubes, and an equal amount of AMPure XP beads was added, followed by incubation at room temperature for 5 min after vortexing. The mixture was placed on a magnetic station for 5 min, and the supernatant was removed. The magnetic beads were washed twice with 200 μL of 80% ethanol. Finally, DNA was eluted with 20 μL of 10 mM Tris-HCl (pH 7.5).

### 3. 16S rRNA gene amplicon sequencing

Library preparation using a two-step PCR amplification protocol has been reported in our previous publication [14]. In this study, we compared two purification methods: magnetic beads-based purification and exonuclease after the first PCR step. Briefly, the V4 region of bacterial 16S rRNA gene was amplified with 515f and 806rB primers (forward primer: 5′- TCG TCG GCA GCG TCA GAT GTG TAT AAG AGA CAG- [3–6-mer Ns] – GTG YCA GCM GCC GCG GTA A -3′; reverse primer: 5′- GTC TCG TGG GCT CGG AGA TGT GTA TAA GAG ACA G [3–6-mer Ns] - GGA CTA CNV GGG TWT CTA AT -3′) [34,48]. Each sample (1 μL of 10-fold diluted DNA) was amplified in a 10 μL reaction volume containing 0.2 U KOD FX Neo DNA polymerase (TOYOBO Co., Ltd., Osaka, Japan), 2 × PCR buffer (TOYOBO), 0.4 mM dNTPs (TOYOBO), 0.2 μM forward and reverse primers, and 1 μM blocking primers (mPNA and pPNA, PNA BIO, Inc., Newbury Park, CA). PCR was performed using the following specifications: 94°C for 2 min followed by 35 cycles at 98°C for 10 s, 78°C for 10 s, 55°C for 30 s, 68°C for 50 s, and a final extension at 68°C for 5 min (ramp rate = 1°C/s). The PCR products were then purified by two separate methods (See 3-1 and 3-2). The second PCR was carried out with the following primers: forward primer: 5′- AAT GAT ACG GCG ACC ACC GAG ATC TAC AC - [8-mer index] - TCG TCG GCA GCG TC -3′, and reverse primer: 5′- CAA GCA GAA GAC GGC ATA CGA GAT - [8-mer index] - GTC TCG TGG GCT CGG -3′ [22]. Each sample (0.8 μL of purified product from the first PCR) was amplified in a 10 μL reaction volume containing 0.2 U KOD FX Neo DNA polymerase (TOYOBO), 2 × PCR buffer (TOYOBO), 0.4 mM dNTPs (TOYOBO), 0.3 μM forward and reverse primers, and 1 μM blocking primers (mPNA and pPNA). PCR was performed as follows: 94°C for 2 min, followed by 8 cycles at 98°C for 10 s, 78°C for 10 s, 55°C for 30 s, 68°C for 50 s, and a final extension at 68°C for 5 min (ramp rate = 1°C/s). Following amplification, PCR products for each sample were cleaned and size-selected using AMPure XP beads and washed twice with 80% ethanol. The libraries were eluted from the pellet with 10 µL of 10 mM Tris-HCl pH 7.5, quantified with a microplate photometer (Infinite 200 PRO M Nano^+^, TECAN Japan Co., Ltd.), and pooled into a single library in equal molar quantities. The pooled library was sequenced on an Illumina MiSeq platform using a 2× 300-bp MiSeq Reagent Nano Kit v2 (Illumina, CA, USA).

#### 3-1. AMPure XP beads method for PCR purification

A solution containing AMPure XP beads (10 μL) was added to 10 μL of product obtained from the first PCR, and the mixture was incubated at room temperature for 5 min after mixing by vortexing. The mixture was then placed on a magnetic station for 5 min, and the supernatant was subsequently removed. The magnetic beads were washed twice with 200 μL of 80% ethanol. The purified sample was eluted from the beads by incubation with 10 μL of 10 mM Tris-HCl (pH 7.5).

#### 3-2. Exonuclease method for PCR purification

Two μL of ExoSAP-IT Express (Thermo Fisher Scientific, Cat #75001.1.EA) was added to 5 μL of the product obtained from the first PCR, and the mixture was incubated at 37°C for 4 min, followed by 80°C for 1 min.

### 4. Bioinformatics

Bioinformatics and statistical analyses were carried out using the Quantitative Insights Into Microbial Ecology 2 program (QIIME 2, ver. 2020.6.0, https://qiime2.org/) installed through a docker [49]. The raw paired-end FASTQ files were imported into the QIIME2 program and demultiplexed using a native plugin. Thereafter, the Cutadapt plugin was processed primer-trimmed. The Divisive Amplicon Denoising Algorithm 2 (DADA2) plugin in QIIME2 was used for quality filtering. The demultiplexed FASTQ file was trimmed, de-noised, the chimera was removed, and the data were merged [50]. We applied the parameter with truncation length of 220 for both forward and reverse reads. Taxonomic groups were assigned identity with the Naive Bayes q2-feature-classifier trained using the 515F/806R region from 99% operational taxonomic units (OTUs) from the SILVA 138 rRNA database [51,52]. Contaminating archaeal, eukaryotic, mitochondrial, and chloroplast sequences were filtered out of the resulting feature table. After taxonomic assignment of amplicon sequence variants (ASVs), the remaining representative sequences were aligned with MAFFT and used for phylogenetic reconstruction in IQ-TREE multicore version 2.0.3 [53]. The sampling depth parameter was set to 1,181, which was chosen based on the number of sequences in the sample containing lowest number of sequences (**Additional file 1: Figure S3**). Finally, diversity indices such as Shannon diversity, Faith phylogenetic diversity, Bray-Curtis distance, and weighted UniFrac distance were calculated using the QIIME2 diversity plugin. The resulting data were exported as a BIOM table and imported to the LDA Effect Size (LEfSe) algorithm to determine the differences in biomarkers [54]. The LEfSe was performed with the following parameters: non-parametric factorial Kruskal-Wallis test, pairwise Wilcoxon test (*P* < 0.05), and LDA > 2.0. Rank-Abundance Dominance (RAD) analysis was performed using the R package *RADanalysis* ver. 0.5.5 [55]. These RAD curves display logarithmic species abundances against rank order using the minimum richness (*R* = 40) to normalize the complete ASV table (**Additional file 2: Table S1**).

## Availability of data and materials

The reported DNA sequence data are available in the DDBJ Sequence Read Archive under the accession number DRA011499.

## Abbreviations

Amplicon-seq: amplicon sequencing
TE: Tris-EDTA
LBB: Lysate Binding Buffer
ISO_AMP: Isopropanol method for DNA extraction and AMPure XP beads method for PCR purification
ISO_EXO: Isopropanol method for DNA extraction and exonuclease method for PCR purification
AMP_AMP: AMPure XP beads method for DNA extraction and AMPure XP beads method for PCR purification
AMP_EXO: AMPure XP beads method for DNA extraction and exonuclease method for PCR purification
ASV: Amplicon Sequencing Variant
PCoA: Principal Coordinate Analysis
LEfSe: Linear Discriminant Analysis Effect Size

## Acknowledgements

We thank Ms. Akiko Tsuruta for her technical assistance.

## Author information

Kie Kumaishi and Erika Usui contributed equally to this work.

## Contributions

YI designed the project; KK and EU performed most of the experiments and data analyses; KS and SK participate in bioinformatics analyses; TS, YT, HT, SS, MN, AT, KM, YY, HT, and HI performed filed experiment and collected samples; KK, EU, and YI wrote the manuscript with inputs from all other authors. The authors read and approved the final manuscript.

## Ethics declarations

Ethics approval and consent to participate

Not applicable.

## Consent for publication

Not applicable.

## Supplementary information

**Additional file 1: Figure S1-S3**.

**Figure S1**. Agarose gel electrophoresis of genomic DNA extracted with different methods.

**Figure S2**. Normalized Rank Abundance Dominance (NRAD) plots obtained from different methods.

**Figure S3**. Rarefaction curve to determine the read number for the analysis.

**Additional file 2: Additional file 2: Table S1**.

Rarefied abundance matrices of the observed ASVs

