## Additional file 1 for "Simple amplicon sequencing library preparation for plant root microbial community profiling"

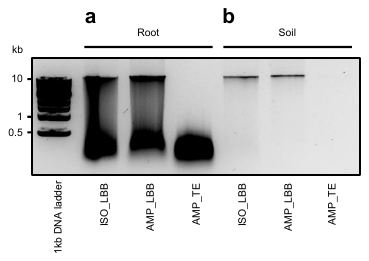

**Figure S1. Agarose gel electrophoresis of genomic DNA extracted with different methods**

The fragmentation status of the extracted DNA from the root (**a**) and soil samples (**b**) are shown.

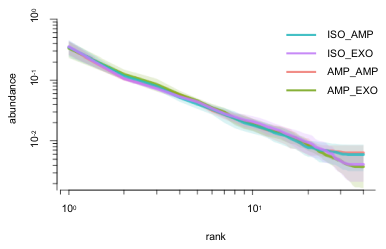

**Figure S2. Normalized Rank Abundance Dominance (NRAD) plots obtained from different methods**

Representative NRAD plots obtained from different methods are shown. The NRADs with stronger tails in the exonuclease method suggest that this method can detect minor bacteria.

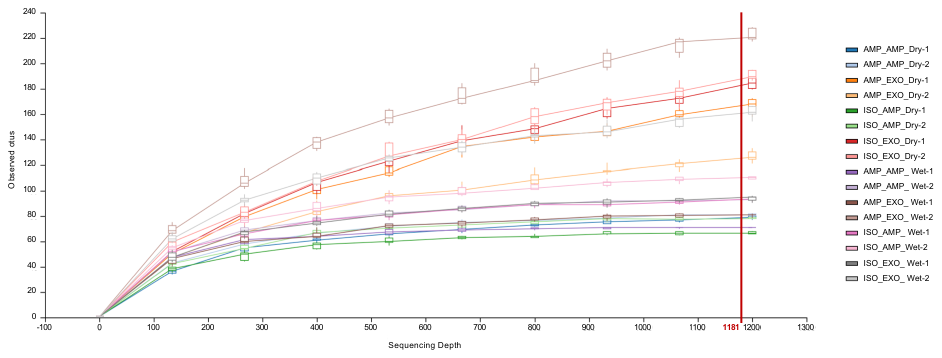

**Figure S3. Rarefaction curve to determine the read number for the analysis**

Rarefaction curves of ASVs across different samples. Based on this result, we determined 1,181 reads as the depth parameter to rescue all samples and capture the differences in the ASVs among samples for the downstream analytical pipeline.
